# Identification and annotation of centromeric hypomethylated regions with Centromere Dip Region (CDR)-Finder

**DOI:** 10.1101/2024.11.01.621587

**Authors:** F. Kumara Mastrorosa, Keisuke K. Oshima, Allison N. Rozanski, William T. Harvey, Evan E. Eichler, Glennis A. Logsdon

## Abstract

Centromeres are chromosomal regions historically understudied with sequencing technologies due to their repetitive nature and short-read mapping limitations. However, recent improvements in long-read sequencing allowed for the investigation of complex regions of the genome at the sequence and epigenetic levels. Here, we present Centromere Dip Region (CDR)-Finder: a tool to identify regions of hypomethylation within the centromeres of high-quality, contiguous genome assemblies. These regions are typically associated with a unique type of chromatin containing the histone H3 variant CENP-A, which marks the location of the kinetochore. CDR-Finder identifies the CDRs in large and short centromeres and generates a BED file indicating the location of the CDRs within the centromere. It also outputs a plot for visualization, validation, and downstream analysis. CDR-Finder is available at https://github.com/EichlerLab/CDR-Finder.

## INTRODUCTION

Centromeres are chromosomal regions essential for the segregation of sister chromatids during cell division. Centromeres are typically composed of near-identical tandem repeats known as α-satellite, with other types of satellites (e.g. β-satellite, γ-satellite, HSat1A, HSat1B, HSat2, and HSat3) found within pericentromeric regions^1,2^. Within the centromere, α-satellite repeats are organized into higher-order repeat (HOR) arrays, which vary in composition and length and constitute the functional unit of the centromeres.

Recently, long-read sequencing technologies such as Pacific Biosciences (PacBio) high-fidelity (HiFi) and Oxford Nanopore Technologies (ONT) long-read sequencing, as well as newly developed genome assembly algorithms^3,4^, have led to the reconstruction of the most complex regions of the human genome, including centromeres^5–8^, telomeres^9^, segmental duplications^10^, and other repetitive regions^9,11^. These^5–9^ and other^12^ studies have also revealed the genetic and epigenetic features of human centromeres. For instance, centromeres are typically hypermethylated throughout the α-satellite HOR array except for a small hypomethylated region called the centromere dip region (CDR)^12^. The CDR was shown to coincide with a unique type of chromatin containing the centromeric histone H3 variant CENP-A^6,7^, which marks the site of the kinetochore. This finding was confirmed with several experimental assays^5–7^.

Here, we present Centromere Dip Region (CDR)-Finder, a tool to identify and annotate the CDRs based on the CpG methylation data obtained from either PacBio HiFi or ONT data. CDR-Finder detects regions of hypomethylation within sequence-resolved centromeric α-satellite HOR arrays, outputs their coordinates and size, and adds annotations based on their characteristics. It can be used to analyze both large and small centromeres, as long as the centromere is fully assembled and sequence-resolved.

## MATERIALS AND METHODS

CDR-Finder requires three input files: a FASTA file of the genome assembly, a BED file containing the coordinates of the region of interest (e.g. the centromere) within the assembly, and a BAM file containing alignments of PacBio HiFi or ONT data with methylation tags to the same genome assembly. CDR-Finder first converts the BAM file to a bedMethyl file using modkit (https://github.com/nanoporetech/modkit), which lists the position and frequency of modified bases for each CpG. Then, it divides the region of interest within the methylBED file into sequential 5-kbp bins and calculates the average CpG methylation frequency for each bin. Bins without an assigned value are excluded. Finally, CDR-Finder runs RepeatMasker^13^ on the region of interest to identify the location of the α-satellite sequences (annotated as “ALR/Alpha”), and it calculates the mean CpG methylation frequency across each bin containing α-satellite.

To identify the CDR(s), our tool first selects bins with a CpG methylation frequency less than the median frequency of all α-satellite sequences in the region of interest. While this frequency can be specified by the user, in our experience, a frequency of 0.34 identifies most CDRs with high precision and recall (described in the example below). Frequencies greater than 0.34 often fail to detect CDRs with shallow hypomethylation, and frequencies less than 0.34 often miscall CDRs due to variation in sequencing coverage. Then, CDR-Finder further refines the bins with a low CpG methylation frequency to those that also have a minimal dip prominence (defined topographically^14^) from the median. In our experience, a minimal dip prominence of 0.30 often removes low-confidence calls when the methylation levels are uniformly low across a subregion. Finally, CDR-Finder evaluates the boundaries of each candidate CDR by calculating the mean CpG methylation frequency and then extending each call boundary to the mean CpG methylation frequency +/-a specific value. In our experience, the mean minus one standard deviation is usually sufficient to capture the entire CDR in each call.

## USAGE AND EXAMPLES

CDR-Finder is organized as a configurable Snakemake pipeline. To run it, the user should first clone the repository from https://github.com/EichlerLab/CDR-Finder. Then, the user should modify the configuration file, config.yaml, to specify the sample name, the path to sample’s genome assembly, the path to the BED file containing a region(s) of interest, and the path to the BAM file containing the alignment of PacBio HiFi or ONT data with methylation tag to the sample’s genome assembly.

In most cases, default parameters can be maintained, but the sequencing coverage and methylation-calling algorithm will affect the ability of CDR-Finder to detect CDRs accurately. As such, the parameters may need to be adjusted accordingly. While CDR-Finder can run on any sequence containing α-satellite DNA, we recommend that the user run it on an α-satellite HOR array with additional flanking sequence on both sides to ensure that the centromere is completely traversed and to observe the transition in methylation patterns between the centromeric and pericentromeric regions. Since CDR detection is based on α-satellite sequences only, the additional flanking sequence will not affect the results of the analysis. We also recommend that the user verify the accuracy of each call based on the coverage data in the plot generated by CDR-Finder and in the original methyl-reads alignment. The user should exclude calls in regions with low coverage and other potential false positives.

The pipeline can be run using conda or singularity with the following commands: `snakemake -np -- sdm conda -c 4` or `snakemake -np --sdm apptainer conda -c 4`, respectively. It will output a BED file with the coordinates of each CDR call as well as a plot showing the CpG methylation frequency of the region of interest, the overall sequencing coverage and methylated sequencing coverage, and annotation showing the location of the CDR call and the sequence composition of the region (**Figure 1**).

**Figure 1.**
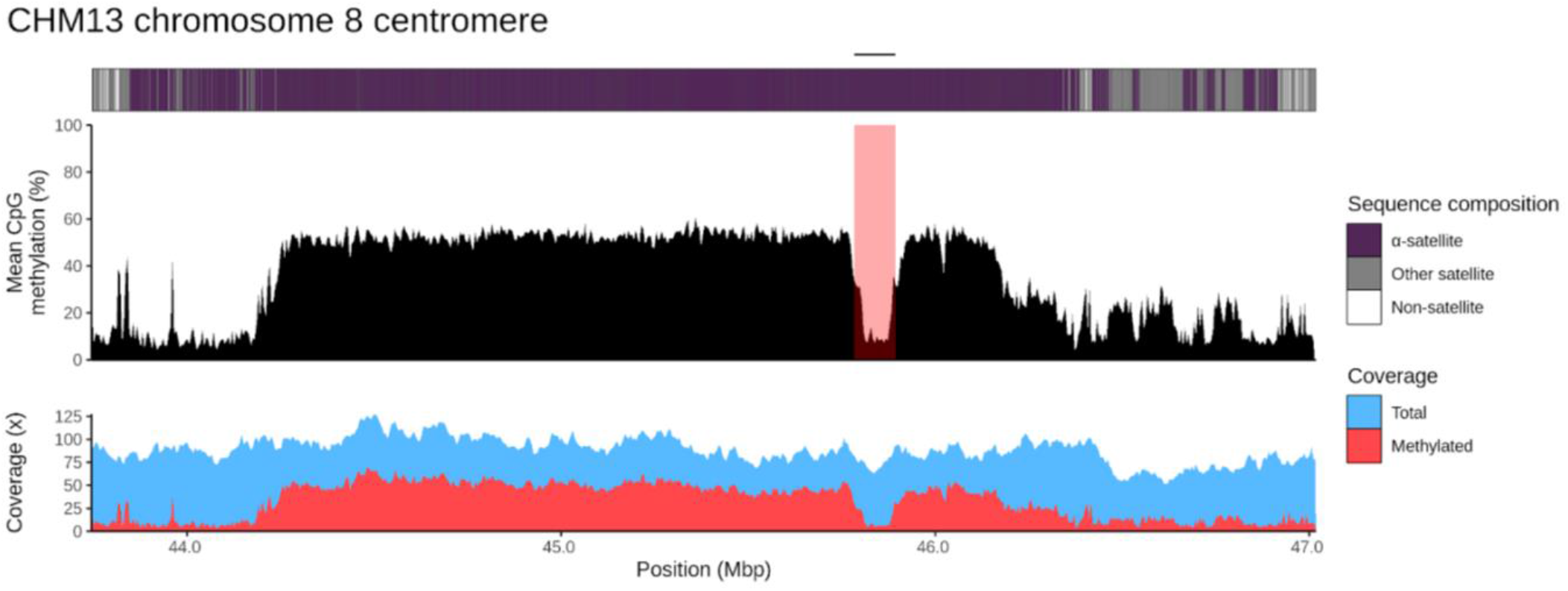
Detection of a CDR in the T2T-CHM13 chromosome 8 centromere with CDR-Finder. CDR-Finder generates a plot with the annotation of the region (top), mean CpG methylation frequency (middle), and the corresponding read coverage (both total and methylated reads) across the region (bottom). The CDR is highlighted in red and indicated with a black bar on top.

Below, we show an example of a CDR call for the T2T-CHM13 chromosome 8 centromere^6^ characterized by CDR-Finder (**Figure 1**). We used the original chromosome 8 centromere α-satellite HOR array coordinates with ∼500 kbp additional sequence on both sides as a target region (chr8:43,746,447-47,020,471 in the T2T-CHM13 v2.0 genome^9^) as well as the original ONT data generated from the same genome. CDR-Finder called only one CDR, as expected (chr8:45,789,626-45,899,626, size = 110 kbp). The same T2T-CHM13 chromosome 8 CDR was originally described with a similar size (73 kbp)^6^. However, on that occasion, CpG methylation was detected with a different method, and the CDR was considered as the region with the lowest CpG methylation levels.

To evaluate CDR-Finder’s ability to detect CDRs in diverse human centromeres, we ran the tool on 200 completely and accurately assembled centromeres from 15 randomly selected samples from the Human Genome Structural Variation Consortium^15^ with default parameters. We manually inspected all calls, counting the number of CDRs that were correctly called, partially called (where the CDR could be extended on one or both sides), incorrectly called, or not called. Our test showed that 98.7% (443/449) of the CDRs were correctly called, indicating a precision of 0.99, and only 43 CDRs were not called (8.7%), indicating a recall of 0.91. In 28 cases (6.3%), the CDR(s) were only partially called, which could be corrected by tweaking the parameters for the specific case. The six erroneous calls could be easily excluded with a visual inspection of the CDR-Finder plots (**Supplementary Table 1)**.

## CONCLUSION

We developed CDR-Finder, a user-friendly method to identify CDRs in complete centromeres. The tool analyzes CpG methylation data from both ONT and PacBio HiFi data and outputs the coordinates of the CDR calls and a plot with related read coverage and highlighted CDR windows. In our experience, both PacBio HiFi and ONT data perform similarly with this tool^16^. However, the user can modify the parameters based on specific cases and quality of the data available. We provide an extensive explanation of the parameters with test cases and common issues on the CDR-Finder GitHub page (https://github.com/EichlerLab/CDR-Finder).

## ACKNOWLEDGMENTS

This research was supported, in part, by funding from the National Institutes of Health (NIH) R01HG010169 (E.E.E.) and R00GM147352 (G.A.L.). E.E.E. is an investigator of the Howard Hughes Medical Institute. This article is subject to HHMI’s Open Access to Publications policy. HHMI lab heads have previously granted a nonexclusive CC BY 4.0 license to the public and a sublicensable license to HHMI in their research articles. Pursuant to those licenses, the author-accepted manuscript of this article can be made freely available under a CC BY 4.0 license immediately upon publication.

## CONFLICTS OF INTERESTS STATEMENT

E.E.E. is a scientific advisory board (SAB) member of Variant Bio, Inc. The other authors declare no competing interests.

## DATA AVAILABILITY

The T2T-CHM13 v2.0 genome assembly, PacBio HiFi data, and ONT data are available at: https://github.com/marbl/CHM13. The HGSVC genome assemblies, PacBio HiFi data, and ONT data are available at: https://ftp.1000genomes.ebi.ac.uk/vol1/ftp/data_collections/HGSVC3/release. The ONT data used in our tests were basecalled with Guppy v6.3.7, which detects methylated cytosines during basecalling, and were aligned to the T2T-CHM13 v2.0 reference genome using winnowmap2^17^. The following command was used for ONT read alignment and processing: `winnowmap -W CHM13_repetitive_k15.txt -y --eqx -ax map-ont -s 4000 -t {threads} -I 10g {ref.fasta} {reads.fastq} | samtools view -u -F 2308 - | samtools sort -o {output.bam} –`.

## Notes

https://github.com/EichlerLab/CDR-Finder

